# ICOS costimulation is indispensable for the differentiation of T follicular regulatory cells

**DOI:** 10.1101/2021.09.14.460384

**Authors:** Vincent Panneton, Barbara C. Mindt, Yasser Bouklouch, Antoine Bouchard, Saba Mohammaei, Jinsam Chang, Mariko Witalis, Joanna Li, Albert Stancescu, John E. Bradley, Troy D. Randall, Jörg H. Fritz, Woong-Kyung Suh

## Abstract

ICOS is a T cell costimulatory receptor critical for Tfh cell generation and function. However, the role of ICOS in Tfr cell differentiation remains unclear. Using Foxp3-Cre-mediated ICOS knockout (ICOS FC) mice, we show that ICOS deficiency in Treg-lineage cells drastically reduces the number of Tfr cells during GC reactions but has a minimal impact on conventional Treg cells. Single-cell transcriptome analysis of Foxp3^+^ cells at an early stage of the GC reaction suggests that ICOS normally inhibits *Klf2* expression to promote follicular features including *Bcl6* upregulation. Further, ICOS costimulation promotes nuclear localization of NFAT2, a known driver of CXCR5 expression. Notably, ICOS FC mice had an unaltered overall GC B cell output but showed signs of expanded autoreactive B cells along with elevated autoantibody titers. Thus, our study demonstrates that ICOS costimulation is critical for Tfr cell differentiation and highlights the importance of Tfr cells in maintaining humoral immune tolerance during GC reactions.

## Introduction

High affinity class-switched antibodies are essential for immune responses against pathogens. These antibodies arise from germinal centers (GCs) where T follicular helper (Tfh) cells facilitate the transition of GC B cells into antibody-secreting plasma cells (PCs) (Crotty, 2011; Victora and Nussenzweig, 2012). Tfh cells are defined by combined expression of their master transcription factor Bcl6 along with CXCR5, PD-1 and ICOS (Fazilleau et al., 2009; Johnston et al., 2009; Nurieva et al., 2009; Yu et al., 2009). Tfh cells mediate their helper functions through costimulation by CD40L and ICOS along with the production of the cytokines IL-4 and IL-21 (Bryant et al., 2007; Crotty, 2014; Reinhardt et al., 2009). Because dysregulation of Tfh cells and GC reactions can lead to humoral autoimmunity, they must be tightly controlled (Linterman et al., 2009).

T follicular regulatory (Tfr) cells are a subset of CD4^+^ Foxp3^+^ regulatory T (Treg) cells found in and around germinal centers (Chung et al., 2011; Linterman et al., 2011; Sayin et al., 2018; Wollenberg et al., 2011). Like Tfh cells, they express the chemokine receptor CXCR5 which is required for their migration towards B cell follicles (Chung et al., 2011; Linterman et al., 2011; Wollenberg et al., 2011). The transcription factor NFAT2 was recently shown to be required for CXCR5 upregulation by Tfrs, possibly to overcome BLIMP-1-mediated CXCR5 downregulation (Oestreich et al., 2012; Shaffer et al., 2002; Vaeth et al., 2014). There are no known lineage-defining factors specific to Tfr cells, although they require concomitant expression of Foxp3 and Bcl6 (Hou et al., 2019; Wu et al., 2016). A significant proportion of Tfr cells originate from thymic Tregs and possess a T cell receptor (TCR) repertoire skewed towards self-antigens (Chung et al., 2011; Linterman et al., 2011; Maceiras et al., 2017; Wollenberg et al., 2011). Under specific conditions, induced Tregs have displayed the capacity to differentiate into Tfr cells that can be specific for the immunizing antigen (Aloulou et al., 2016). Strong IL-2 signaling was shown to inhibit Tfr differentiation which is more akin to Tfh cells rather than Tregs (Botta et al., 2017). While the *in vivo* role of Tfr cells has been controversial, they display suppressive abilities on T cell proliferation, antibody secretion and cytokine production *in vitro* (Sage et al., 2014; Sage et al., 2013). Several Tfr depletion models have been studied to understand their functions *in vivo*. Initially, adoptive transfer or mixed bone marrow (BM) chimera experiments showed that Tfr reduction had varying effects on GC responses, possibly due to unintended side effects such as impaired Treg function (Chung et al., 2011; Linterman et al., 2011; Wollenberg et al., 2011). More recently, Foxp3-specific deletion of Bcl6-expressing (Bcl6 FC) or CXCR5-expressing cells (Tfr-deleter) allowed for a more precise assessment of *in vivo* roles for Tfr cells (Clement et al., 2019; Fu et al., 2018; Laidlaw et al., 2017; Wu et al., 2016; Xie and Dent, 2018). Results from these studies collectively suggest two roles for Tfr cells: suppression of autoantibody production and modulation of antibody responses more suggestive of “helper” functions (Sage and Sharpe, 2020).

The inducible costimulator (ICOS) is a member of the CD28 superfamily and is known to be expressed by activated T cells (Hutloff et al., 1999). ICOS was previously shown to be essential for the formation of Tfh cells and maintenance of Bcl6 expression (Bossaller et al., 2006; Gigoux et al., 2009; Leavenworth et al., 2015). In both mice and humans, ICOS null mutations cause severe defects in GC reactions and antibody production due to the lack of Tfh cells (Dong et al., 2001; Grimbacher et al., 2003; McAdam et al., 2001; Tafuri et al., 2001). Some ICOS-deficient patients develop autoimmune symptoms such as rheumatoid arthritis and autoimmune neutropenia, suggesting a potential role for ICOS in Treg/Tfr compartments (Takahashi et al., 2009; Warnatz et al., 2006). Indeed, ICOS deficiency in mice led to reduced Tfr cell numbers, although the underlying mechanisms have not been carefully analyzed (Sage et al., 2013; Zhang et al., 2018).

In this study, we used *Icos^fl/fl^ Foxp3-Cre* (ICOS FC) mice to evaluate the role of ICOS signaling in Treg and Tfr cells during GC reactions. Foxp3-specific loss of ICOS led to a significant decrease in Tfr populations after protein immunization or viral infection without affecting Treg cell numbers. Examination of antibody responses revealed significantly lowered IgG2b titers at steady state or after immune challenge along with increased anti-nuclear autoantibodies in ICOS FC mice. Single-cell transcriptomics and biochemical analyses suggest that ICOS may enhance the Treg to Tfr transition through KLF2 and NFAT2 regulation. Overall, our findings indicate that the major role of ICOS in regulatory T cell compartments during GC reactions is to control Tfr differentiation and highlight the importance of Tfr cells in preventing autoantibody generation.

## RESULTS

### ICOS is required for Tfr cell generation during GC reactions against protein antigens

To assess the role of ICOS in Treg lineage cells, we used a Foxp3-Cre system which allows for the specific abrogation of ICOS expression in all Treg and Tfr cells. Throughout this study, we used *Icos^+/+^ Foxp3-Cre^+^* controls (ICOS WT) for *Icos^fl/fl^ Foxp3-Cre^+^* mice (ICOS FC). First, we analyzed splenocytes 12 days after immunization with NP-OVA/alum by flow cytometry and subdivided the CD4^+^ Foxp3^+^ regulatory T cell compartment into Treg (CD4^+^ Foxp3^+^ CXCR5^-^ PD-1^-^), PD-1^-^ Tfr (CD4^+^ Foxp3^+^ CXCR5^+^ PD-1^-^) and PD-1^+^ Tfr (CD4^+^ Foxp3^+^ CXCR5^-^ PD-1^+^) subsets (Fig. 1 A). ICOS was expressed in both Tfh (CD4^+^ Foxp3^-^ CXCR5^+^ PD-1^+^) and Treg/Tfr cells, with PD-1^+^ Tfr cells showing the highest surface levels (Fig. S1 A).

**Figure 1.**
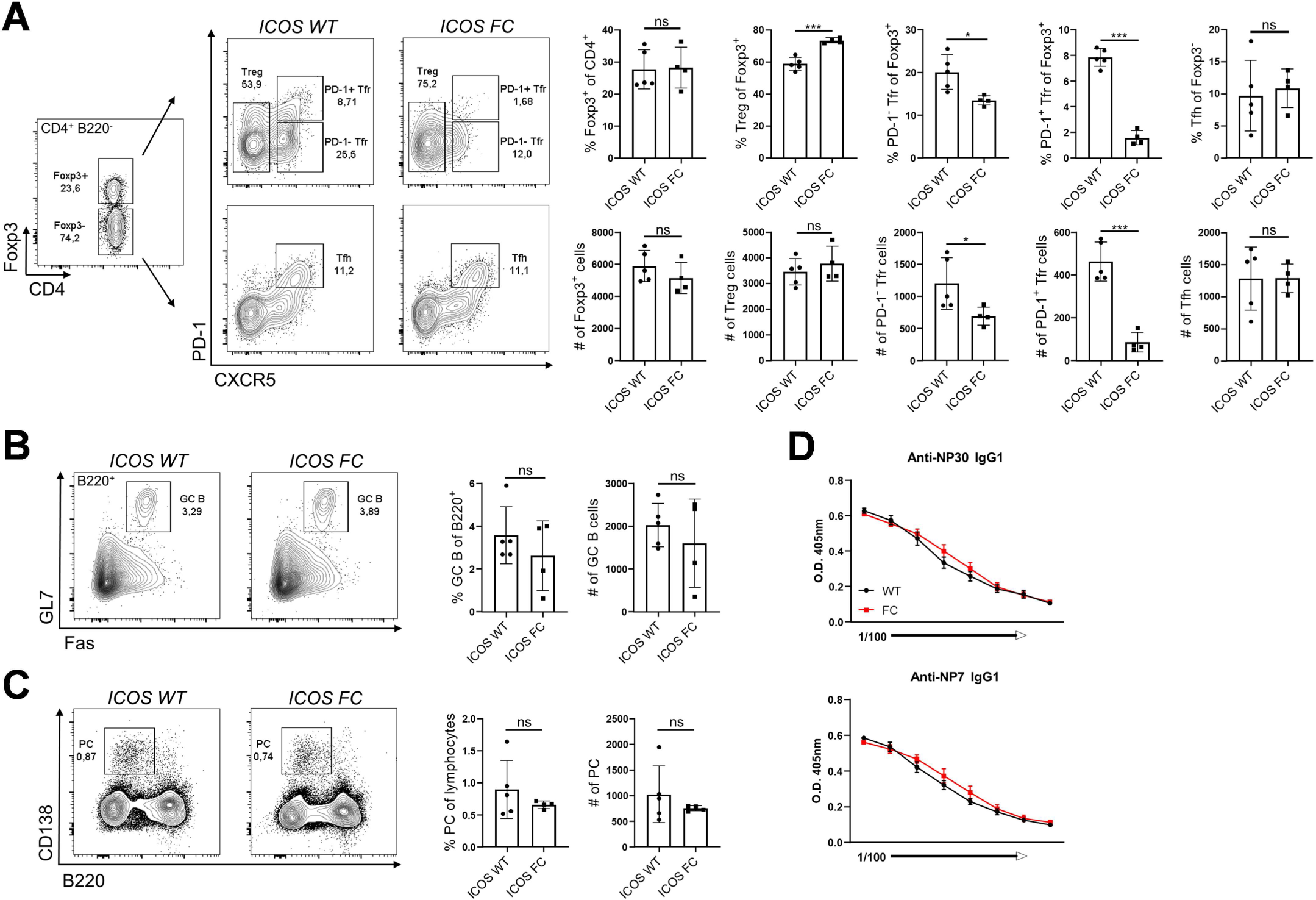
Foxp3-specific ICOS ablation decreases Tfr populations in a protein immunization model. Splenocytes from ICOS WT (n=5) and ICOS FC (n=4) mice were harvested 12 days post-immunization (dpi) with NP-OVA/alum and analyzed by flow cytometry. (A) Treg, Tfr and Tfh cell percentages and numbers were evaluated using a combination of Foxp3, PD-1 and CXCR5 staining. (B) B220^+^ Fas^+^ GL7^+^ GC B cell populations and (C) B220^-^ CD138^+^ plasma cells populations were analyzed from the same splenocyte pool 12 dpi. (D) Total (NP30) and high affinity (NP7) anti-NP IgG1 titers were measured by ELISA using serum from ICOS WT (black, n=5) or ICOS FC mice (red, n=5) obtained 28 dpi. Data shown as mean ± SEM, \**p*<0.05, \*\**p*<0.01, \*\*\**p*<0.001. All data are representative of at least three independent experiments.

We confirmed that ICOS deletion was limited to Foxp3^+^ cells and did not occur in Tfh cells (Fig. S1 B). We observed no change of total Foxp3^+^ populations in ICOS FC mice (Fig. 1 A). However, we found ∼2-fold and ∼4-fold decreases in PD-1^-^ and PD-1^+^ Tfr cell proportions and numbers, respectively. Interestingly, these reduced Tfr populations were balanced by an increased proportion of Treg cells. Further, we observed that Foxp3-specific ICOS abrogation leads to a ∼4-fold reduction of Foxp3^+^ cells within the GC (Fig. S2). Next, we examined Tfh, GC B cells and plasma cells since lack of Tfr cells has been shown to increase these populations in some experimental settings (Chung et al., 2011; Linterman et al., 2011; Wollenberg et al., 2011). We did not observe significant differences in Tfh cell proportion or absolute numbers (Fig. 1 A). We also detected no quantitative differences in GC B cell and plasma cell populations (Fig. 1 B-C). Consistently, NP-specific IgG1 antibodies in serum did not show changes in total (NP30) or high affinity (NP7) titers (Fig. 1 D). Taken together, these results indicate that loss of ICOS in Foxp3^+^ cells leads to a reduced number of Tfr cells with proportionally increased Treg cells, suggesting an impaired Treg-to-Tfr transition.

#### ICOS is required for Tfr generation during anti-viral response

To evaluate the role of ICOS in regulatory T cell compartments during an anti-viral immune response, we infected mice with influenza A virus (IAV). We analyzed splenocytes by flow cytometry 30 days after infection since it has been shown that Tfr generation is delayed due to high levels of IL-2 present in the early stages of viral infection (Botta et al., 2017). We observed an increased proportion of Tregs and decreases in Tfr populations in spleens of ICOS FC mice reminiscent of results from protein immunization experiments (Fig. 2 A). As before, we did not detect significant differences in Tfh populations (Fig. 2 B). Next, we examined the expansion of GC B cell populations that recognize the IAV nucleoprotein using recombinant tetramers (Flu tetramer) (Allie et al., 2019). Interestingly, we observed a trend of increased total GC B cells with a significantly decreased proportion of IAV-specific GC B cells in ICOS FC mice (Fig. 2 C). This results in a significant increase of extraneous GC B cells, some of which could be autoreactive in nature. On the other hand, we did not observe significant differences in splenic plasma cells (Fig. 2 D). However, we found significant decreases of IAV-specific IgG2b serum titers (Fig. 2 E). Consistent with the role of anti-viral antibodies in the overall control of influenza virus (Lam and Baumgarth, 2019), we observed that ICOS FC mice had more severe weight loss 9-10 days post-infection (Fig. S3 A). This reduced anti-viral IgG2b titer was well correlated with a reduction of total IgG2b titers in infected ICOS FC mice (Fig. S3 B). Further, we noticed that uninfected ICOS FC mice had decreased basal levels of IgG2b and IgG1 (Fig. S3 C), suggesting that ICOS-expressing Treg or Tfr cells may work as “helper” T cells for certain antibody isotype switching. Thus, congruent with data from protein immunization experiments, this infection model confirmed the critical role of ICOS for efficient Tfr generation. Further, increases of non-viral specific GC B cells in ICOS FC mice confirms the regulatory role of Tfr cells in shaping GC responses.

**Figure 2.**
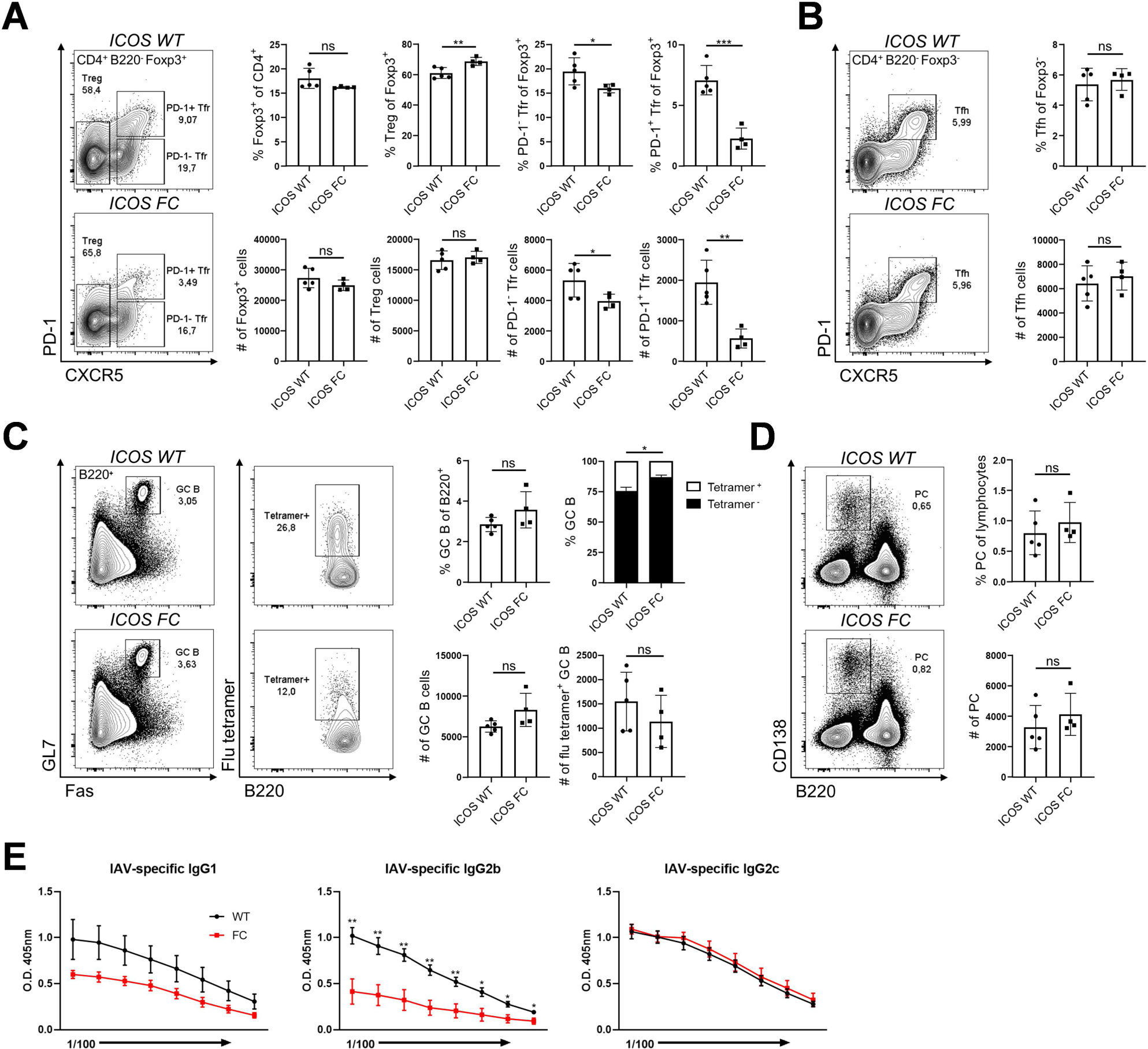
ICOS FC mice display reduced Tfr populations along with increased extraneous GC B cells during anti-viral responses. Splenocytes from ICOS WT (n=5) or ICOS FC (n=4) were harvested 30 days post-infection (dpi) with influenza A virus (IAV) and analyzed by flow cytometry. (A) Foxp3^+^ Treg and Tfr cells were classified using CXCR5 and PD-1 staining. (B) Foxp3^-^ CXCR5^+^ PD-1^+^ Tfh cells were analyzed from the same splenocyte pool. (C) B220^+^ Fas^+^ GL7^+^ GC B cells were harvested from spleens 30 dpi and stained with influenza nucleoprotein-specific tetramers (flu tetramers). (D) B220^-^ CD138^+^ plasma cells were analyzed using the same splenocyte pool. (E) IAV-specific IgG1, IgG2b and IgG2c titers were measured by ELISA using serum samples from ICOS WT (black) and ICOS FC mice (red) obtained 30 dpi. Data shown as mean ± SEM, \**p*<0.05, \*\**p*<0.01, \*\*\**p*<0.001. All data are representative of two independent experiments.

### Treg-specific ICOS deficiency leads to anti-nuclear antibody production

Since there is ongoing cell death and release of autoantigens within the GC, autoreactive GC B cell clones can expand and differentiate into PCs with help from Tfh cells if not restrained (Linterman et al., 2009). Given that Foxp3-specific ICOS ablation results in reduced numbers of Tfr cells which are thought to suppress self-reactive antibody production, we investigated whether ICOS FC mice displayed signs of autoimmunity. We did not observe immune infiltration in the kidneys, lungs, spleen, pancreas, and salivary glands of 5-month-old ICOS FC mice (Fig. 3 A). Next, we used HEp-2 slides to look for the presence of anti-nuclear antibodies (ANAs) which are a hallmark of autoimmunity (Fig. 3 B) (Castro and Gourley, 2010). We did not detect significant spontaneous increases of ANAs in serum samples of 6-month-old ICOS FC mice. However, we found that both single NP-OVA/alum immunization and secondary challenge with the same antigen resulted in significantly higher ANAs in ICOS FC mice. Additionally, ICOS FC mice infected with influenza A virus presented similar increases in ANA levels. These results suggest that immunization or infection augments adventitious generation of autoantibodies which is normally suppressed by Tfr cells.

**Figure 3.**
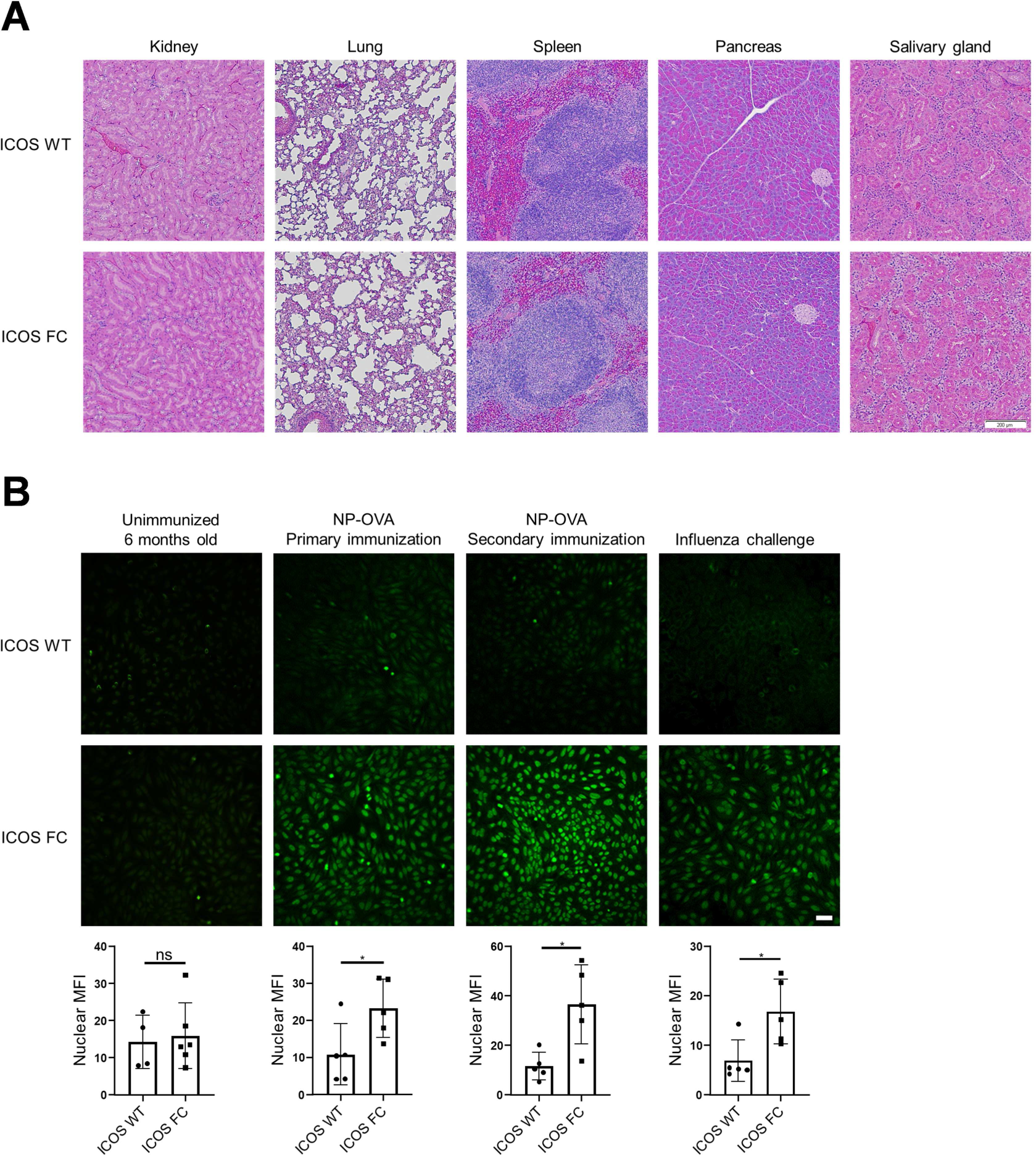
ICOS FC mice produce increased anti-nuclear antibodies following immune challenge. (A) Representative images of H&E-stained tissue sections from 5-month-old ICOS WT (n=3) and ICOS FC mice (n=3). Scale bar 200 µM. (B) Anti-nuclear antibodies were detected by staining HEp-2 slides with serum samples collected from mice after the following treatments. Unimmunized ICOS WT (n=4) and ICOS FC (n=6) mice at 6 months of age. Primary immunization of ICOS WT (n=5) and ICOS FC (n=5) mice with NP-OVA/alum (serum harvested 44dpi). Secondary NP-OVA/alum injection 30 days after the primary injection (serum harvested 44 days post secondary challenge). IAV infection of ICOS WT (n=5) and ICOS FC (n=5) mice (serum harvested 30 dpi). Scale bar 50µM. Nuclear fluorescence intensity was quantified using ImageJ. Data shown as mean ± SEM, \**p*<0.05. All data are representative of two independent experiments.

### ICOS-deficient Treg cells show impaired transition to Tfr cells

To better understand the role of ICOS in Tfr cell differentiation, we performed single-cell transcriptome analysis of CD4^+^ Foxp3^+^ splenocytes sorted from ICOS WT and ICOS FC mice immunized with NP-OVA/alum. To collect cells in a dynamic Treg-to-Tfr transition stage, we prepared samples 6 days post-immunization, a timepoint where Tfr cells begin to appear (Botta et al., 2017; Sage et al., 2013). After sorting, we added back sorted conventional CD4^+^ Foxp3^-^ T cells (∼10% of total) into the sorted CD4^+^ Foxp3^+^ T cell pool to provide a reference point (Fig. S4 A, cluster 4). We used Tfr-defining genes (*Cxcr5, Pdcd1, Foxp3, Bcl6*) to calculate a “Tfr identity score” and selected three clusters that are predicted to contain Tfr precursors and mature Tfr cells (Fig. S4 B, clusters 3, 5, 8 black arrows). When compared amongst each other, these cells formed three distinct clusters with a spectrum of Tfr identity score (Fig. 4 A, clusters 1, 2, 3). Pseudotime trajectory analysis revealed a progressive differentiation from cluster 1 towards cluster 3 (Fig. 4 B). Interestingly, we observed that ICOS FC mice presented a 3-fold increase of cells in cluster 2 and a 3-fold decrease in cluster 3 (Fig. 4 A). This suggests that Tregs could be halted in their transition to Tfr cells due to loss of ICOS expression. To substantiate this idea, we compared gene expression profiles of the three clusters (Fig. 4 C, left). Cluster 3 had the highest levels of key Tfr signature genes (*Cxcr5, Pdcd1, Bcl6*), but reduced expression of typical Treg signature genes such as *Foxp3, Il2ra* (CD25), *Ctla4* and *Tnfrsf18* (GITR) when compared to clusters 1 and 2. However, when compared with CD4^+^ Foxp3^-^ conventional T cells, cluster 3 still mostly maintained higher levels of these Treg signature genes (Fig 4 C, right). Recent studies have identified CD25 downregulation as a key event in Tfr differentiation (Botta et al., 2017; Wing et al., 2017).

**Figure 4.**
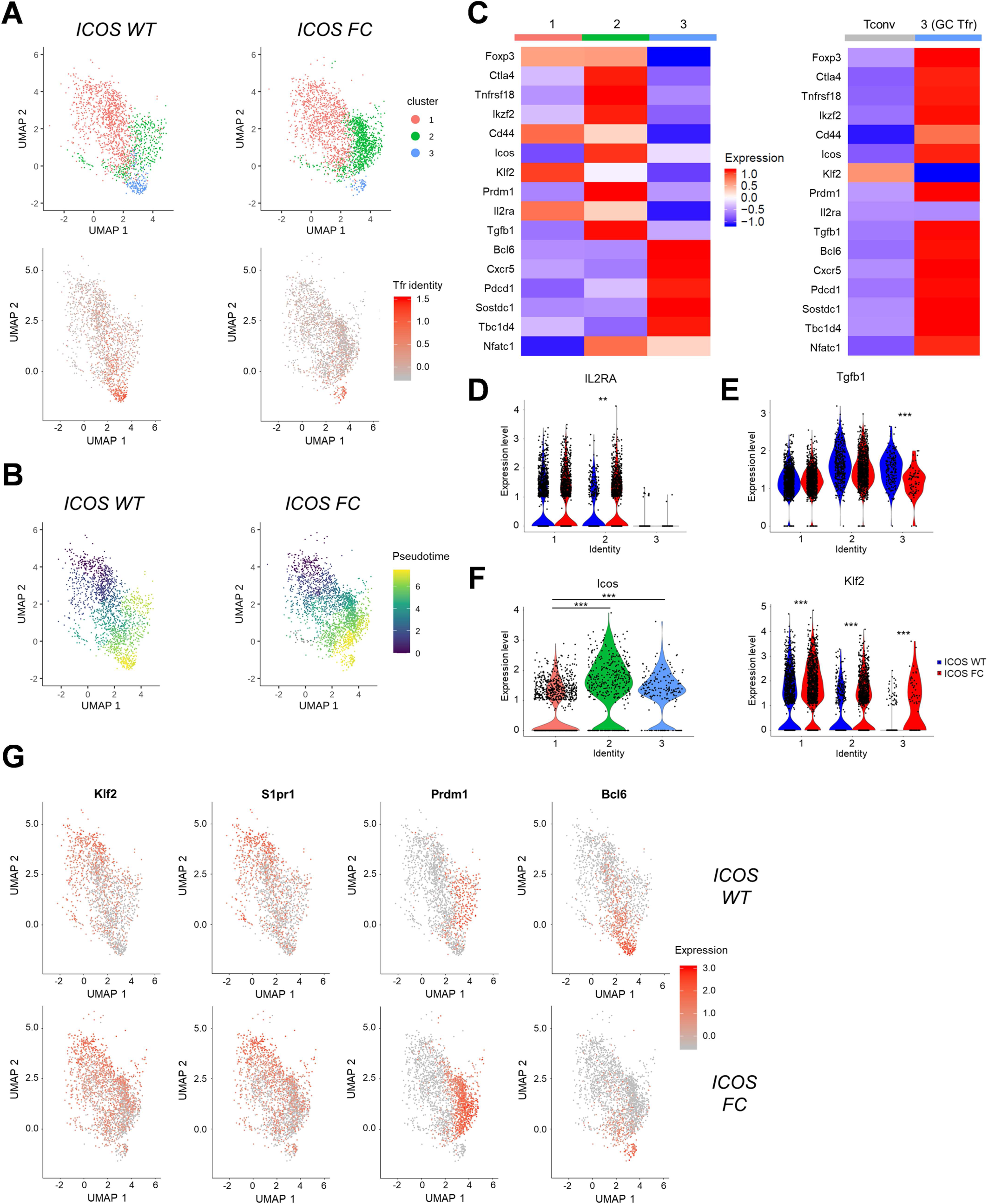
ICOS-deficient Treg cells show impaired Treg to Tfr transition. Single-cell transcriptomes of FACS-sorted CD4^+^ Foxp3^+^ splenocytes from an ICOS WT or ICOS FC mouse harvested 6 days after protein immunization. (A) Selection and sub-clustering of Foxp3^+^ cells based on positive Tfr identity scores. (B) Pseudotime analysis showing the differentiation trajectory of selected Foxp3^+^ splenocytes. (C) Mean expression of regulatory and follicular genes by the indicated subpopulation. (D) *Il2ra* (E) *Tgfb1*, (F) *Icos* and *Klf2* violin plots showing the gene expression levels subdivided by cluster identities defined in (A). (G) Feature plots of *Klf2, S1pr1, Prdm1* and *Bcl6* expression. Each dot represents one cell. \**p*<0.05, \*\**p*<0.01, \*\*\**p*<0.001.

Similarly, we noticed that CD25 expression is inversely correlated with the levels of CXCR5 and PD-1 in Tfr cells, consistent with a previous report by Wing *et al*. (Wing et al., 2017) (Fig. S5 A-B). Congruent with CD25 protein expression levels, *Il2ra* gene expression was significantly dampened in cluster 3 Tfr cells compared to those in clusters 1 and 2 (Fig. 4 D). Interestingly, cluster 2 cells from ICOS FC mice displayed significantly higher *Il2ra* levels suggesting that ICOS could be involved in CD25 downregulation. ICOS deletion also resulted in significantly elevated CD25 protein expression in certain Tfr subsets (Fig. S5 B). Apart from CD25, we noticed a significant decrease of *Tgfb1* expression in ICOS FC cluster 3 cells (Fig. 4 E). This could explain reduced IgG2b titers observed in ICOS FC mice since TGF-β1 is a known class switch factor for this isotype (McIntyre et al., 1993).

Another key event in the Treg-to-Tfr transition is CXCR5 upregulation (Chung et al., 2011; Linterman et al., 2011; Vaeth et al., 2014; Wollenberg et al., 2011). ICOS was previously shown to regulate CXCR5 by suppressing KLF2 expression in Tfh cells (Weber et al., 2015). This mechanism may operate in Treg/Tfr cells since we observed that *Icos* and *Klf2* gene expression is inversely correlated and that all three clusters had higher *Klf2* expression in ICOS FC mice (Fig. 4 F). KLF2 was also shown to dampen Tfh differentiation by increasing S1PR1 and BLIMP-1 expression levels (Lee et al., 2015). We observed matching expression patterns of *Klf2, S1pr1 and Prdm1* with opposed *Bcl6* expression in all clusters (Fig. 4 G). Further, ICOS FC mice showed an accumulation of *Klf2^+^ S1pr1^+^ Prdm1^+^* cells in cluster 2 with reduced *Bcl6* expression in clusters 2 and 3. Thus, these results suggest that ICOS is required for a few key steps in the Treg to Tfr transition and that failure of these processes seem to lead to an accumulation of putative Tfr precursors.

### ICOS ablation causes decreased NFAT2 activation and impaired CXCR5 expression in Tregs

To test the potential role of ICOS in upregulating CXCR5 expression, we further analyzed our single-cell transcriptome data. We found that clusters 1 and 2 from ICOS FC mice had lowered proportions of *Cxcr5*^+^ cells along with a diminished average *Cxcr5* expression (Fig. 5 A). Next, we investigated the potential impacts of ICOS signaling on NFAT2 (product of *Nfatc1* gene), a transcription factor known to directly bind to the promoter region of C*xcr5* (Vaeth et al., 2014). Importantly, we and others have previously shown that ICOS can potentiate TCR-mediated calcium flux, a key factor in NFAT activation (Hogan et al., 2003; Leconte et al., 2016). Consistently, we found that the average expression levels of known NFAT target genes (Table S1) was significantly decreased in ICOS FC cluster 2 cells when compared to ICOS WT control (Fig. 5 B, top). Since Tfr precursor-like cells in cluster 2 also express the highest levels of *Nfatc1* (Fig. 4 C), we tested whether *Nfatc1* expression level was reduced in ICOS FC cluster 2 cells. However, we did not find significant differences in the expression level of the *Nfatc1* gene itself (Fig. 5 B, bottom). Nonetheless, we noticed that the protein levels of ICOS and NFAT2 trended higher in Tfr cells compared to Treg cells, suggesting a potential role of ICOS-NFAT2 in the Treg-to-Tfr transition (Fig. 5 C). Consistent with our single-cell transcriptomics data, we found that both PD-1^-^ and PD-1^+^ Tfr subsets from infected ICOS FC mice displayed significantly decreased CXCR5 expression (Fig. 5 D). Congruent with the unaltered *Nfatc1* mRNA levels in ICOS FC Tfr populations, we found that NFAT2 protein expression levels were not decreased in ICOS-deficient Tfr subsets (Fig. 5 E).

**Figure 5.**
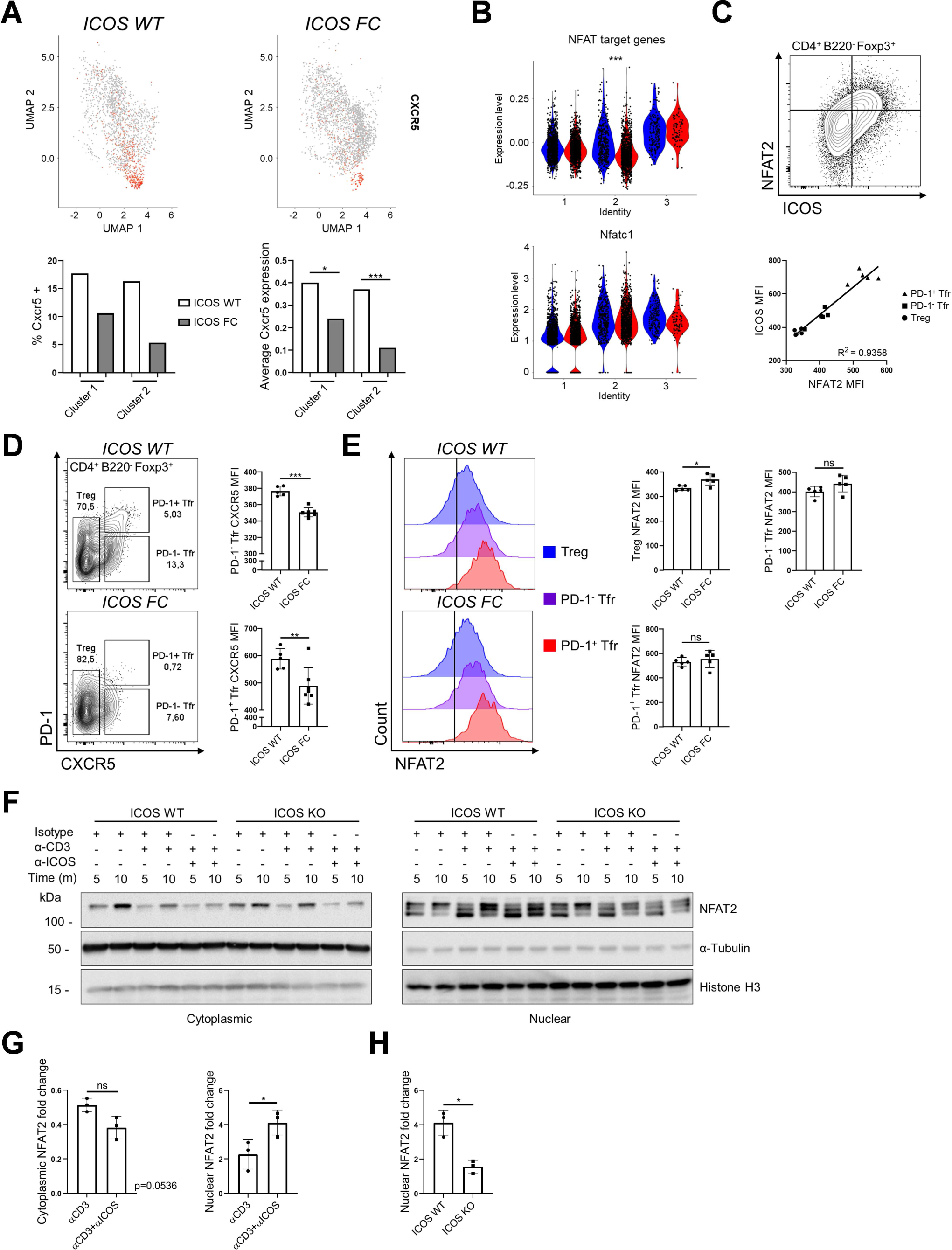
ICOS-NFAT2 signaling may regulate CXCR5 expression in regulatory T cells. (A) Feature plot of *Cxcr5* gene expression (top) and quantification of *Cxcr5*^+^ cells along with average *Cxcr5* expression in clusters 1 & 2 cells (bottom). (B) Average expression of NFAT target genes (top) and expression of *Nfatc1* (bottom) subdivided by cluster identity. (C) Splenocytes from ICOS WT (n=5) and ICOS FC mice (n=7) were analyzed 30 dpi with IAV. ICOS and NFAT2 expression was compared in Treg, PD-1^-^ Tfr and PD-1^+^ Tfr cells coming from ICOS WT mice. (D) MFI values for CXCR5 in PD-1^-^ Tfr and PD-1^+^ Tfr cells from the same splenocyte pool. (E) NFAT2 MFI values in ICOS WT vs. ICOS FC Treg, PD-1^-^ Tfr and PD-1^+^ Tfr cells were measured using the same splenocyte pool. (F) ICOS WT and ICOS germline KO CD4^+^ T cells were isolated from spleens by magnetic sorting and cultured for 2 days with α-CD3/CD28 stimulation. Cells were then restimulated for the indicated times with combinations of isotype control, α-CD3 and α-ICOS antibodies after which cytoplasmic and nuclear fractions were extracted and analyzed for phospho-NFAT2 by Western blot. (G) Bar graph representing normalized cytoplasmic vs. nuclear NFAT2 fold change over isotype control of the indicated samples at the 10min timepoint. Only the bottom NFAT2 band was quantified to represent hypophosphorylated nuclear NFAT2. (H) Normalized nuclear NFAT2 fold change of α-CD3/α-ICOS stimulated samples from ICOS WT vs. KO cells at 10 min. Data shown as mean ± SEM, \**p*<0.05, \*\**p*<0.01, \*\*\**p*<0.001. All data are representative of three independent experiments.

Because we observed that ICOS deletion led to decreased expression of NFAT target genes but not NFAT2 itself, we investigated other regulatory mechanisms. It has been established that NFAT2 activity depends on its dynamic nuclear-cytoplasmic shuttling controlled by its phosphorylation status (Hogan et al., 2003). To evaluate if ICOS can regulate this process, we expanded purified splenic CD4^+^ T cells *in vitro* and acutely restimulated them through the TCR (CD3) with or without ICOS costimulation. Cytoplasmic and nuclear NFAT2 proteins of varying phosphorylation status were then harvested through subcellular fractionation and quantified by Western blot (Fig. 5 F). Without restimulation, 5-10 min incubation at 37°C led to the increase of hyperphosphorylated (slower migrating) NFAT2 species in the cytoplasm with concomitant disappearance of hypophosphorylated (faster migrating) NFAT2 species in the nucleus (lanes 1 and 2 in cytoplasmic and nuclear fractions). This is presumably due to temperature-induced shifts in the activity of NFAT phosphatases and kinases (Hogan et al., 2003). As predicted, CD3 ligation increased NFAT2 nuclear levels (lanes 3 and 4). Importantly, combined CD3/ICOS stimulation significantly augmented nuclear NFAT2 levels at 10 min when compared with CD3 stimulation alone (Fig. 5 G). This increase was dependent on ICOS since it was abolished when ICOS KO CD4^+^ T cells were used (Fig. 5 H). Taken together, our biochemical data shows that ICOS costimulation can potentiate TCR-driven NFAT2 activation in CD4^+^ T cells. Thus, we propose that ICOS could act upstream of the NFAT2-CXCR5 signaling axis known to be one of the key mechanisms for early Tfr differentiation (Vaeth et al., 2014).

## Discussion

In this study, we identified ICOS as a critical costimulatory receptor for Tfr differentiation upon immune challenges. We show that Foxp3-specific ICOS ablation results in altered gene expression patterns at the single-cell level leading to an accumulation of Tfr precursor-like cells and a substantial reduction of the fully differentiated “GC Tfr” population. Mice with impaired Tfr differentiation showed an increase of extraneous GC B cells without increases of total GC B cell or Tfh cell numbers. ICOS FC mice also showed an elevated incidence of autoantibody production after GC reactions, presumably due to an expansion of autoreactive B cells. In contrast, total and virus-specific IgG2b antibody titers under steady-state and upon influenza infection were diminished in ICOS FC mice. Our single-cell data strongly suggests that ICOS-mediated downregulation of KLF2 plays a key role in shaping chemokine receptor expression and balancing BLIMP-1-Bcl6 levels in developing Tfr cells. Our biochemical analysis demonstrates that ICOS signaling can also augment NFAT2 nuclear localization, potentially counterbalancing the negative impacts of BLIMP-1 on CXCR5 expression (Oestreich et al., 2012; Vaeth et al., 2014).

Our single-cell RNA transcriptome analysis of Foxp3^+^ cells from ICOS WT and ICOS FC mice indicates that dynamic changes in gene expression patterns drive the Treg-to-Tfr transition. Trajectory analysis reveals that the CD25^+^ BLIMP-1^+^ activated Treg subset progressively gains follicular features such as CXCR5 and Bcl6. Both single-cell transcriptomics and flow cytometry data demonstrate that Bcl6 and CXCR5 levels are highest in CD25^-^ “GC-Tfr” cells, the main Foxp3^+^ cells shown to be found within the GC (Wing et al., 2017). During this Treg-to-Tfr transition, ICOS seems to utilize a mechanism that has been shown to be critical for the differentiation and maintenance of Tfh cells – timely downregulation of KLF2 (Lee et al., 2015). As such, Tfr precursor cells reduced levels of *Klf2* and its main target genes *S1pr1* and *Prdm1* along the predicted Tfr trajectory. In contrast, *Cxcr5* and *Bcl6* expression levels were progressively elevated in cells that have dampened *Klf2* target genes, presumably due to the lack of BLIMP-1-mediated suppression of Bcl6. Importantly, this progression towards “GC-Tfr” was halted at the *Klf2*^hi^ *Prdm1*^hi^ stage in ICOS-deficient Treg cells. Therefore, we propose that ICOS-mediated KLF2 downregulation is a key molecular event that initiates follicular T cell programing in Tfr cells.

In addition to KLF2, ICOS may utilize NFAT2-dependent pathways to support CXCR5 expression. Although NFAT2 is highly expressed in both Tfh and Tfr cells, abrogation of NFAT2 expression in T cells was shown to cause more pronounced defects in Tfr generation as opposed to Tfh differentiation due to compromised CXCR5 expression (Vaeth et al., 2014). This observation fits well with the idea that developing Tfr cells need higher concentrations of nuclear NFAT2 to overcome elevated levels of BLIMP-1 (known repressor of *Cxcr5* gene (Oestreich et al., 2012)) that are present in Tfr precursor cells. Our biochemical data indicate that ICOS ligation augments the amount of nuclear NFAT2 in TCR-activated CD4^+^ T cells. Based on these, we speculate that ICOS costimulation reinforces CXCR5 expression in early Tfr populations leading to the establishment of “GC-Tfr” differentiation.

While we showed that ICOS signaling can maintain NFAT2 in the nucleus, the mechanism remains unclear. Nuclear transport of NFAT family members occurs through their dephosphorylation by calcineurin, a Ca^2+^ dependent phosphatase (Hogan et al., 2003). We have previously shown that ICOS signaling can potentiate TCR-induced intracellular Ca^2+^ flux, although we did not determine if this resulted in increased NFAT activity (Leconte et al., 2016). Conversely, nuclear export of NFAT2 is triggered by phosphorylation through several kinases including GSK3β (Beals et al., 1997). In turn, GSK3β activity can be inhibited by Akt-mediated phosphorylation of residue Ser9 (Yoeli-Lerner et al., 2009). We and others have shown that ICOS stimulation can increase PI3K/Akt signaling, specifically through its Y^181^MFM cytoplasmic tail motif (Arimura et al., 2002; Gigoux et al., 2009; Parry et al., 2003). Thus, we suggest that ICOS could maintain NFAT2 nuclear localization by increasing its import through enhanced Ca^2+^ signaling and/or decrease its export by inhibiting GSK3β.

The biological roles of Tfr cells during GC reactions remain ill-defined. Tfr depletion studies using Bcl6 FC mice have shown small decreases of antibodies specific for the immunizing antigen (Wu et al., 2016; Xie et al., 2020; Xie et al., 2019). One study also showed spontaneous increases of autoantibodies and multi-organ lymphocytic infiltration in aged mice (Fu et al., 2018). Along the same line, high levels of anti-nuclear antibodies were generated in Bcl6 FC mice after influenza infection (Botta et al., 2017). ICOS FC mice do not display age-related autoantibodies or lymphocytic infiltrations, suggesting that the impact of ICOS-deficiency on Treg cells is weaker than that of Bcl6-deficiency. In this context, Bcl6-deficient Treg cells (as opposed to bona-fide Tfr cells) could have contributed to some of these phenotypes considering recent reports that Bcl6-deficient Tregs have compromised suppressive functions in other immune settings (Li et al., 2020; Sawant et al., 2012; Sawant et al., 2015). ICOS deficiency was shown to impair the suppressive ability of Tregs in asthma and type 1 diabetes murine models, but no defects in GC reactions were reported (Busse et al., 2012; Kornete et al., 2012). Combined with a ∼4-fold reduction in the Tfr number, it seems likely that humoral immune defects in ICOS FC mice are mainly due to reduced Tfr numbers. Another potential role for Tfr cells is to promote the generation of antigen-specific antibodies. Bcl6 FC mice produce reduced amounts of IgE and IgG1 in certain immunization protocols (Xie et al., 2020; Xie et al., 2019). We found that both basal IgG2b (in unimmunized mice) and anti-influenza IgG2b titers are lower in ICOS FC mice. This correlated well with reduced amounts of TGF-β1 mRNA in the GC-Tfr population. However, further work is required to clarify the mechanisms and biological implications of this finding.

In sum, we showed that ICOS is critically important for Tfr differentiation. ICOS-mediated downregulation of KLF2 and its target genes can shape the Bcl6-driven Tfr programming whereas an ICOS-NFAT2-CXCR5 signaling axis may reinforce CXCR5 expression during Tfr differentiation. Our data supports the view that the main role of Tfr cells is to suppress the expansion of self-reactive GC B cells during GC reactions, and we believe that our ICOS FC mouse provides a complementary model to dissect Tfr differentiation and function.

## Materials and methods

### Mice and animal procedures

C57BL/6 and *Foxp3^YFP-Cre^* mice (Jax 016959) (Rubtsov et al., 2008) were purchased from the Jackson Laboratory. ICOS conditional knockout mice were generated in C57BL/6 background as previously described (Panneton et al., 2018). ICOS germline knockout mice have been backcrossed onto C57BL/6 background for more than ten generations (Tafuri et al., 2001). All mice were housed in the Institut de Recherches Cliniques de Montréal animal care facility under specific pathogen-free conditions. Animal experiments were performed in accordance with animal use protocols approved by the Institut de Recherches Cliniques de Montréal Animal Care Committee. We used 8–12-week-old male mice for experiments involving Foxp3-Cre-mediated gene deletion unless specified otherwise. For protein immunization, mice were injected intraperitoneally with 100µg of 4-Hydroxy-3-nitrophenylacetyl hapten-17 (NP17)-OVA (Biosearch Technologies, 1µg/mL) mixed with Imject Alum (ThermoFisher) in a 1:1 ratio. For viral infections, mice were infected intranasally with a sublethal dose of influenza A virus H1N1 (strain A/Puerto Rico/8/34 (PR8), 10 PFU/20g body weight).

### Flow cytometry

For analysis, single-cell suspensions were prepared by mechanical disruption of spleens unless specified otherwise. Viability was assessed by staining 1x10^8^ cells/mL with fixable viability dye eFluor 780 (ThermoFisher) for 20 mins at 4°C. Fc receptors were blocked using anti-CD16/CD32 (BioXCell). For intracellular staining, cells were fixed and permeabilized using the Transcription Factor Staining Buffer Set (ThermoFisher). Surface or intracellular staining was performed at 1x10^8^ cells/mL for 20 mins at 4°C. The following antibodies were used. BD Biosciences: anti-CD4 BUV395 (GK1.5), anti-PD-1 BV421 (RMP1-30), anti-CD95 PE-Cy7 (Jo2), anti-Bcl6 PE (K112-91). ThermoFisher: anti-B220 PerCP-eFluor 710 (RA3-6B2), anti-CXCR5 Biotin (SPRCL5), Streptavidin PE-Cy7, anti-ICOS FITC (7E.17G9), anti-Foxp3 APC (FJK-16s), anti-B220 eFluor 450 (RA3-6B2), anti-CD25 PE (PC61.5). BioLegend: anti-NFATc1 PE (7A6), anti-GL7 FITC (GL7). To identify influenza-specific B cells, we used tetramerized recombinant nucleoproteins conjugated with APC or PE (Flu tetramer) provided by Dr. Troy Randall (Allie et al., 2019). Data was acquired using a BD LSR Fortessa and analyzed using FlowJo v10 (BD Biosciences).

### ELISA

Serum samples were obtained from blood collected from the submandibular vein at the indicated timepoints. Plates were coated with either goat anti-mouse IgG (SouthernBiotech), NP30-BSA and NP7-BSA (Biosearch Technologies), or heat-inactivated influenza A viruses overnight at 4°C. Serum samples underwent a 2-fold serial dilution starting from the indicated initial dilution. Bound antibodies were detected using alkaline phosphatase-conjugated anti-IgG1/2b/2c/3, IgM or IgA and *p-*nitrophenyl phosphate substrate (SouthernBiotech). The reaction was stopped by adding 1.5 N NaOH solution and optical density was measured at 405 nm.

### Histology

Organs were dissected and fixed in 10% neutral buffered formalin for 12 h at 4°C. Organs were then washed in 1x PBS, embedded in paraffin, and cut into 5 µM sections. Slides were stained with H&E to examine immune cell infiltration of organs. For immunofluorescence staining of Tfr cells, spleens were fixed for 2 h at 4°C in 2 mL of 4% paraformaldehyde (Millipore Sigma) followed by an overnight incubation at 4°C in 2 mL of 30% sucrose (Millipore Sigma). Next, five consecutive 15 min washes at 4°C in 2L of 30% sucrose were performed. Sucrose was washed out and spleens were frozen in O.C.T. medium (Tissue-Tek). Then, 10 µM sections were cut and permeabilized for 60 mins at RT with 2% Triton X in PBS (Millipore Sigma). Slides were stained at 4°C overnight with a cocktail of anti-mouse CD4 PE (RM4-5), anti-mouse IgD efluor 450 (11-26), anti-mouse GL7 Alexa Fluor 488 (GL-7), and anti-mouse Foxp3 APC (FJK-16s) (ThermoFisher). Fluorescent signals were visualized using a DM6000 fluorescence microscope (Leica).

### Anti-nuclear antibody assay

Serum samples were obtained from blood collected from the submandibular vein at the indicated timepoints and incubated on Kallestad HEp-2 slides (Bio-Rad) according to the manufacturer’s instructions. Bound antibodies were detected using goat anti-mouse IgG Alexa Fluor 555 (ThermoFisher). Fluorophore signals were visualized using a DMRB fluorescence microscope (Leica). Nuclear MFI was quantified using ImageJ.

### Single-cell RNA sequencing

Splenocytes from *Foxp3^YFP-cre^Icos^+/+^* (ICOS WT) and *Foxp3^YFP-cre^Icos^fl/fl^* (ICOS FC) male mice were isolated 6 dpi with NP-OVA/alum and stained with anti-CD4 Alexa Fluor 647 (GK1.5, BioLegend), anti-TCRβ PE-Cy7 (H57-597, BioLegend) and propidium iodide (ThermoFisher). Live (PI^-^) conventional (YFP^-^) and regulatory (YFP^+^) CD4^+^TCRβ^+^ T cells were sorted with a BD FACS Aria (BD Biosciences) to >95% purity. Sorted conventional and regulatory T cells were mixed in a 1:10 ratio to provide an internal control. A total of 13500 cells from ICOS WT and ICOS FC mice were sent for library preparation. Libraries were generated using the following components from 10x Genomics: Chromium Next GEM Chip G Single Cell kit, Chromium Next GEM Single Cell 3’ GEM, Library & Gel Bead kit v3.1, Chromium i7 Multiplex kit. Sequencing was performed by Genome Québec using a NovaSeq 6000 (Illumina) with a flow cell S1 PE28*91.

#### Reads alignment

Using *Cellranger* 4.0.0 (from *10xGenomics*®), we generated a custom reference genome using the GRCm38.p6 (mm10) assembly procured from *Ensembl* to which we added the *Ires-Yfp-iCre* sequence as described in its design map (Rubtsov et al., 2008). The alignment of the reads was performed using the same software and the resulting expression matrix was loaded into *R* version 3.6.1 (from the R Foundation for Statistical Computing) to conduct analysis.

#### Single cell expression matrix analysis

The expression matrices were stored into an R Seurat object available in the package *Seurat* version 3.0 (Stuart et al., 2019) to ease the analysis. ICOS WT and ICOS FC samples were merged during the filtering phase which consisted of the elimination of any cell that presented more than 10% mitochondrial RNA contamination as well as any cell with less than 200 unique genes expressed. The expression matrix was then log normalized and scaled. We identified the most differentially expressed genes within the samples and proceeded with a dimensional reduction using a principal component analysis (PCA) approach based on the 2000 most variable features. We selected the first 30 most important eigenvectors produced by the PCA to construct a *Shared Nearest Neighbor* (SNN) graph and used *Modularity Optimizer version 1.3.0* (Waltman, 2013) to identify 13 clusters. The cells were projected on a 2D space using a *Uniform Manifold Approximation and Projection UMAP* method (McInnes, 2020). We isolated 3 clusters of interest based on their markers and moved the normalized expression matrix into an R *cell_data_set* object available in the package *Monocle3* version 0.2.3 (Trapnell et al., 2014). Using the dimensionally reduced matrices of expression, a differentiation trajectory was constructed. The cells were then ordered along the trajectory and pseudotime was computed. We further confirmed the consistency of our trajectory analysis using a diffusion map-based approach (available in package *destiny* version 2.0.4) which has proven to be more robust to noise (Angerer et al., 2016). We computed NFAT signaling gene expression score using the *AddModuleScore* function available in the *R* library *Seurat v3.0*. The list of NFAT target genes was established using the PANTHER classification system combined with data from literature and can be found in Supplementary table 1 (Hermann-Kleiter and Baier, 2010; Mi et al., 2021; Mognol et al., 2016; Vaeth et al., 2014).

### CD4^+^ T cell activation and Western blot analysis

CD4^+^ T cells were isolated from spleens and lymph nodes using the EasySep mouse CD4^+^ T cell isolation kit (StemCell Technologies) according to the manufacturer’s instructions. Purified T cells were stimulated from 2 days in complete RPMI 1640 (10% FBS, 1 unit/mL penicillin, 1 µg/mL streptomycin, 55mM β-mercaptoethanol and 10 mM HEPES) with plate-bound anti-CD3 (BioXCell, 3 µg/mL) and soluble anti-CD28 (ThermoFisher, 2 µg/mL). For restimulation, CD4^+^ T cell blasts were incubated for 3 mins at room temperature with the indicated combination of the following antibodies: 1 µg/mL Armenian hamster IgG isotype control (BioXCell), 1 µg/mL anti-CD3e (145-2C11, ThermoFisher) and 2 µg/mL anti-ICOS (C398.4a, BioLegend). Goat anti-hamster IgG (20 µg/mL, SouthernBiotech) was added for crosslinking and cells were immediately incubated for the indicated timepoints in a 37°C water bath. Restimulation was stopped using ice-cold STOP buffer (PBS, 10% FBS, 1 mM Na_3_VO_4_, 1 mM EDTA). Cytoplasmic and nuclear fractions of restimulated T cells were obtained using the NE-PER kit (ThermoFisher) according to the manufacturer’s instructions. Lysates were boiled in Laemmli buffer and samples were run on SDS-PAGE gels. Proteins were transferred to Amersham nitrocellulose membranes (GE Healthcare). Membranes were blocked using 3% BSA in TBS-T. The following antibodies were used for detection according to manufacturer’s instructions: anti-NFAT2 (D15F1, Cell Signaling Technology), anti-α-tubulin (2144, New England Biolabs), anti-Histone H3 (4499, New England Biolabs) and anti-mouse IgG-HRP (Santa Cruz Biotechnology). Detection was performed using Amersham ECL prime kits (GE healthcare) and images were captured using a ChemiDoc imaging system (Bio-Rad). Band quantification was performed using ImageJ and normalized to loading controls.

### Statistical analysis

Data is presented as mean ± SEM unless specified otherwise. For single comparisons, statistical significance was judged using two-tailed Student *t-*tests. For multiple comparisons, the Holm-Sidak *t-*test was used. For single-cell gene expression comparisons between clusters, the Wilcoxon signed-rank test was used. R^2^ values were obtained by linear regression. Statistical significance was judged based on *p* values and is indicated as follows: \**p*<0.05, \*\**p*<0.01, \*\*\**p*<0.001. Analysis was performed using Prism 7 (GraphPad software).

## Data accessibility

Single-cell transcriptome data have been deposited in the GEO database under the accession number GSE164995.

## Author contributions

Vincent Panneton and Woong-Kyung Suh conceived and supervised the study. Vincent Panneton, Barbara C. Mindt, Yasser Bouklouch, Saba Mohammaei, Jinsam Chang, Mariko Witalis, Joanna Li, and Albert Stancescu performed the experiments. Vincent Panneton, Yasser Bouklouch, Antoine Bouchard, and Woong-Kyung Suh analyzed the data. John E. Bradley, Troy D. Randall and Jörg H. Fritz contributed key reagents and resources. Vincent Panneton and Woong-Kyung Suh wrote the manuscript. Vincent Panneton, Barbara C. Mindt, Yasser Bouklouch, Mariko Witalis, Joanna Li, Antoine Bouchard, and Woong-Kyung Suh commented and revised the manuscript.

## Acknowledgments

The authors thank Manon Laprise, Viviane Beaulieu, Stéphanie Lemay, Julie Lord, Éric Massicotte, Dominic Filion, and Simone Terouz for their technical assistance. This work was supported by Canadian Institutes of Health Research (PJT 159526, W-K.S.). The authors declare no competing financial interests.

---

**Supplementary Figure 1.**
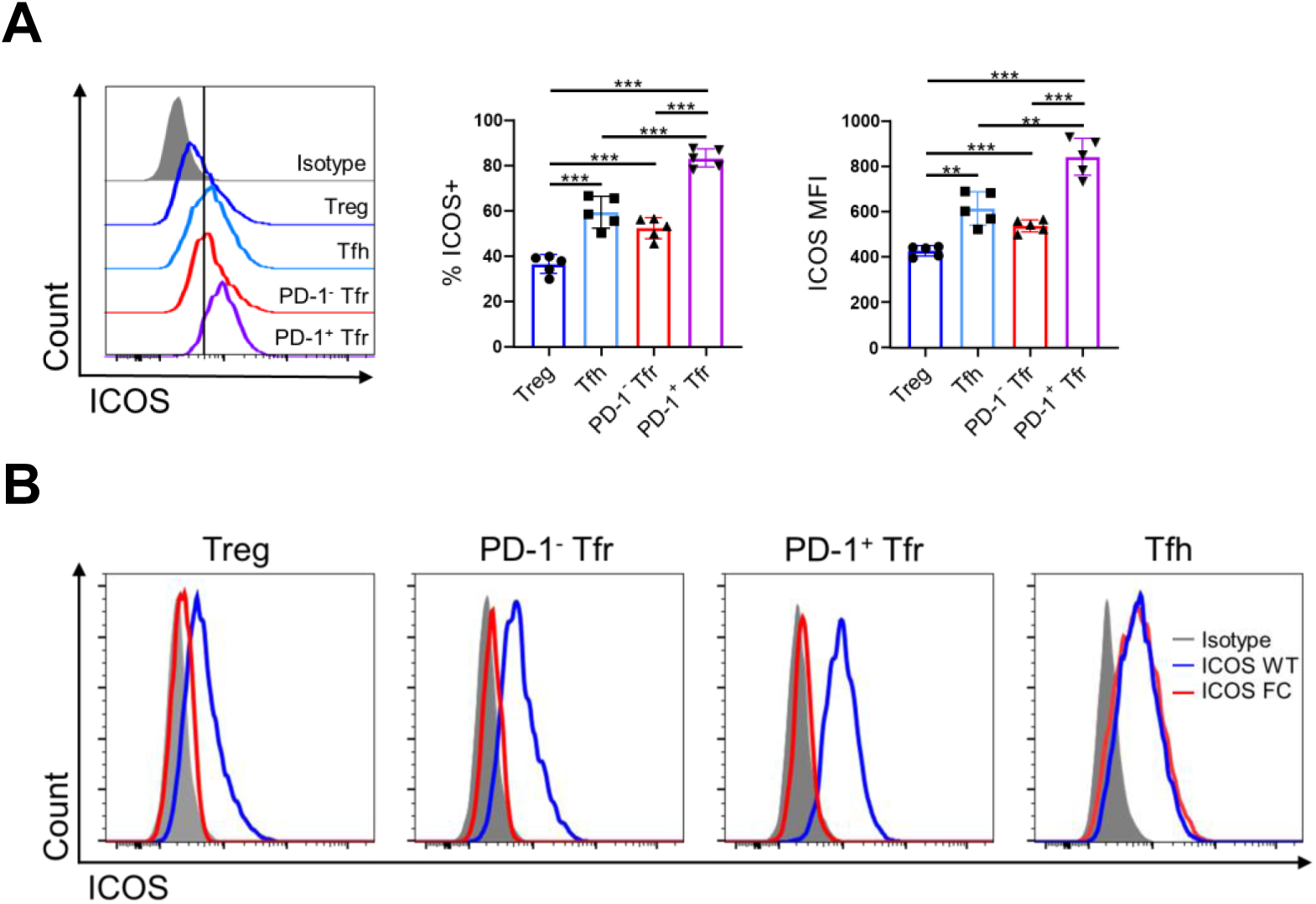
Foxp3-cre mediated ICOS gene deletion is specific to regulatory T cells. (A) Splenocytes from ICOS WT mice (n=5) were analyzed 12 dpi with NP-OVA/alum to measure ICOS expression in the indicated T cell subsets by flow cytometry. (B) Representative histograms of ICOS expression in the indicated T cell subsets in ICOS WT (blue) vs. ICOS FC (red) splenocytes. Data shown as mean ± SEM, \*\**p*<0.01, \*\*\**p*<0.001. All data are representative of three independent experiments.

**Supplementary Figure 2.**
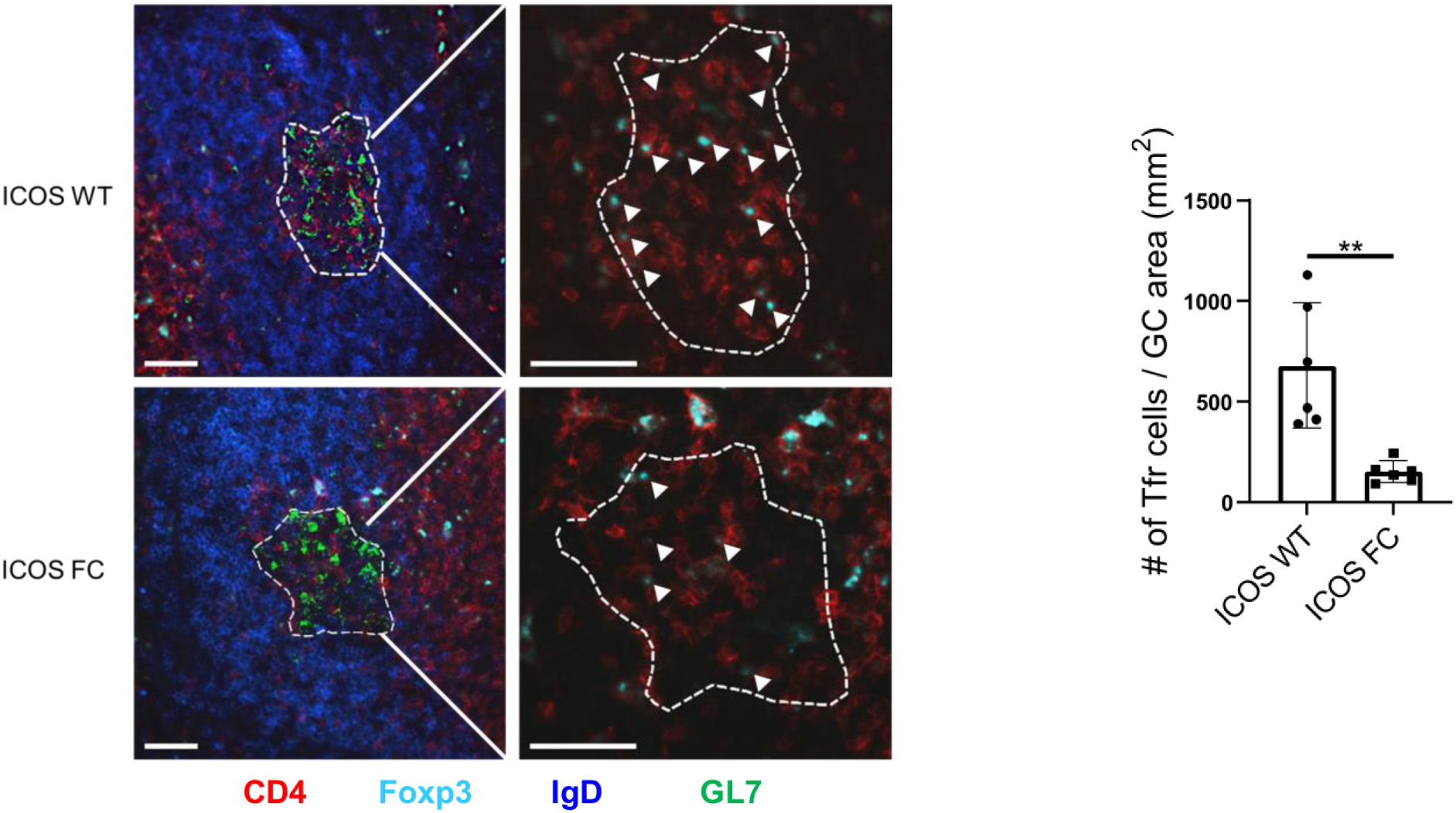
Foxp3-cre mediated ICOS gene deletion reduces GC resident Foxp3+ cell populations Representative images of splenic germinal centers (dotted lines) from ICOS WT or FC mice 12 days post-immunization with NP-OVA/alum. The number of CD4^+^Foxp3^+^ Tfr cells (white arrows) per mm^2^ of GC area was assessed. Scale bar 50 μM. Data shown as mean ± SEM, \**p*<0.05, \*\**p*<0.01. All data are representative of at least two independent experiments.

**Supplementary Figure 3.**
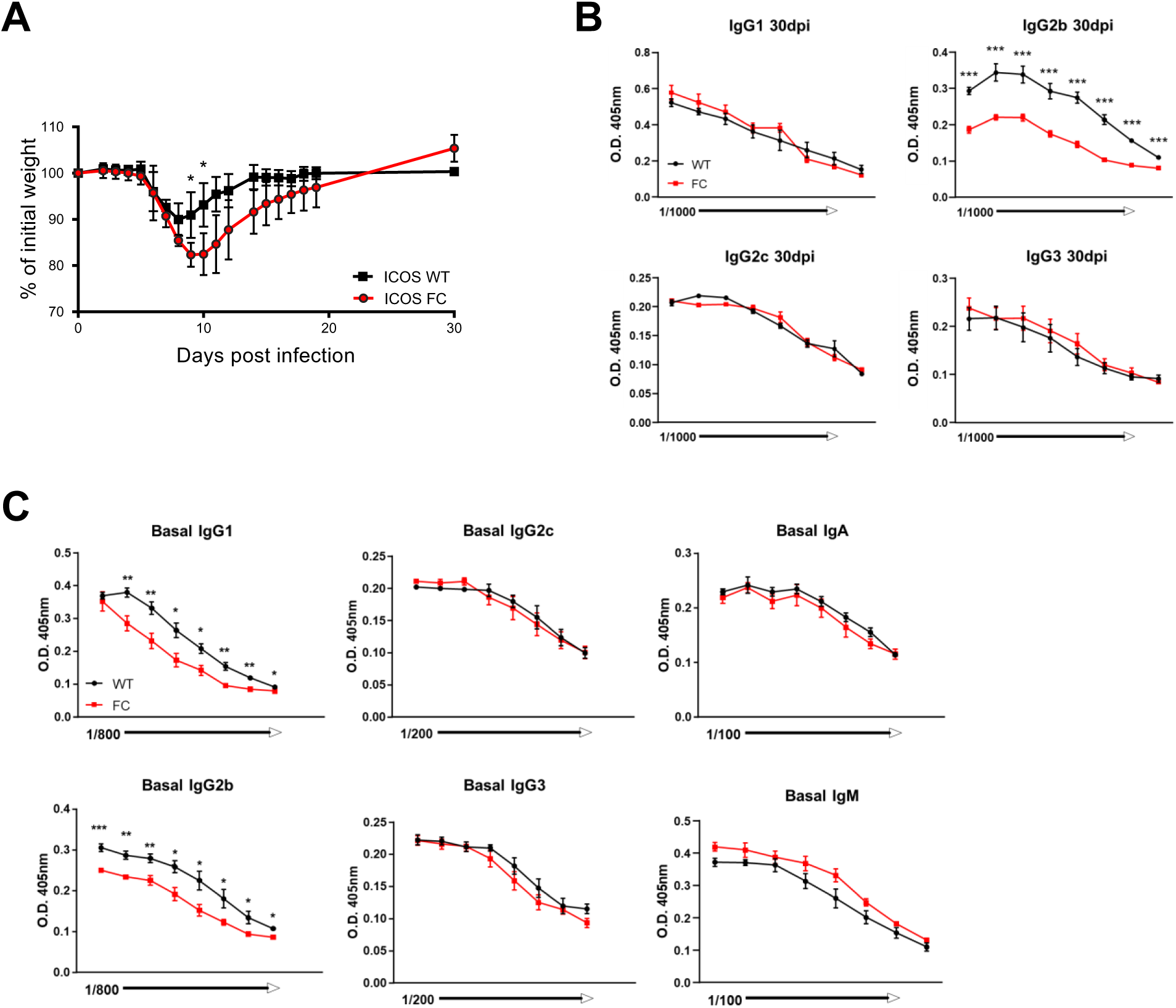
Impaired anti-viral responses in ICOS FC mice. (A) Relative body weight loss over time following influenza A virus (IAV) infection in ICOS WT (n=5) and ICOS FC mice (n=7). (B) Serum samples from ICOS WT (black) and ICOS FC mice (red) was harvested 30 dpi with IAV and total titers of the indicated antibody isotypes were measured by ELISA with 2-fold serial dilutions. (C) Basal antibody titers were measured by ELISA with 2-fold serial dilutions using serum samples from unimmunized ICOS WT (n=6) or ICOS FC mice (n=6). Data shown as mean ± SEM, \**p*<0.05, \*\*\**p*<0.001. All data are representative of two independent experiments.

**Supplementary Figure 4.**
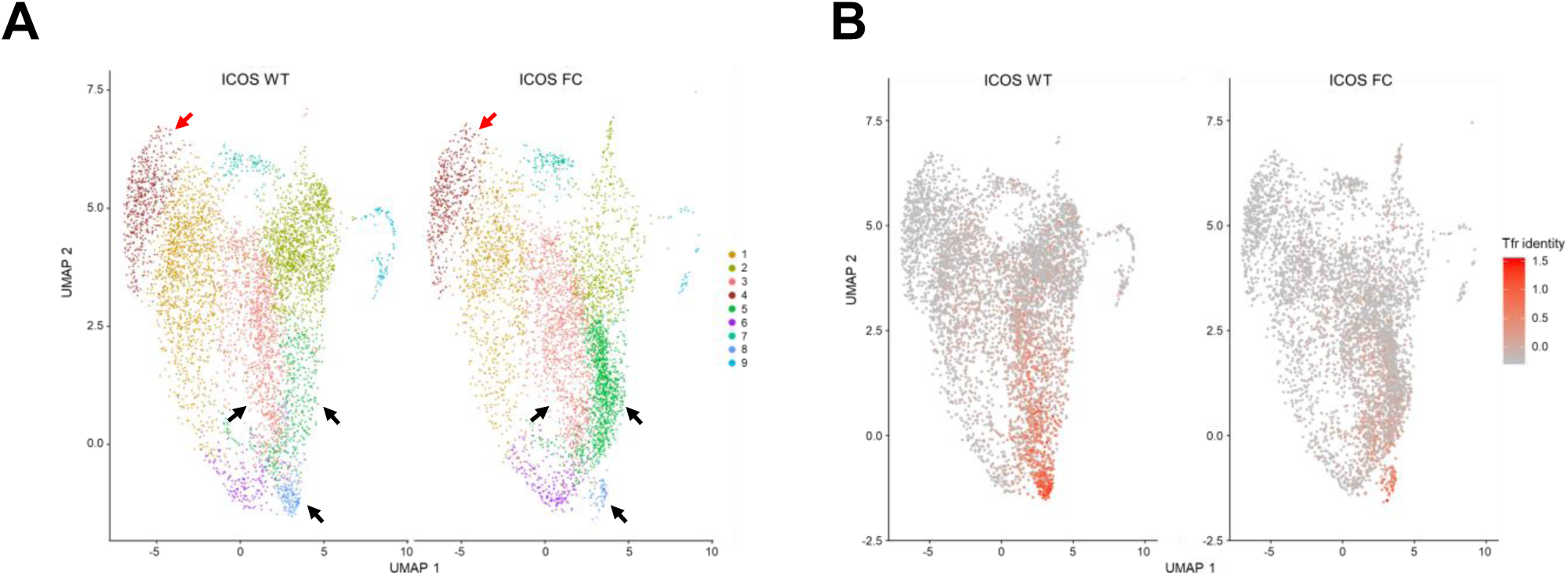
Identification of putative Tfr precursor populations from Treg clusters. (A) UMAP projections of CD4+ Foxp3+ cells isolated from an ICOS WT or ICOS FC mouse 6 dpi with NP-OVA/alum. CD4+ Foxp3-conventional T cells (cluster 4, red arrows) were added back after sorting to provide an internal control. Black arrows represent Tfr-like clusters (3, 5, 8) selected for further analysis. (B) Feature plots of Tfr identity scores based on expression levels of *Foxp3, Cxcr5, Pdcd1* and *Bcl6*.

**Supplementary Figure 5.**
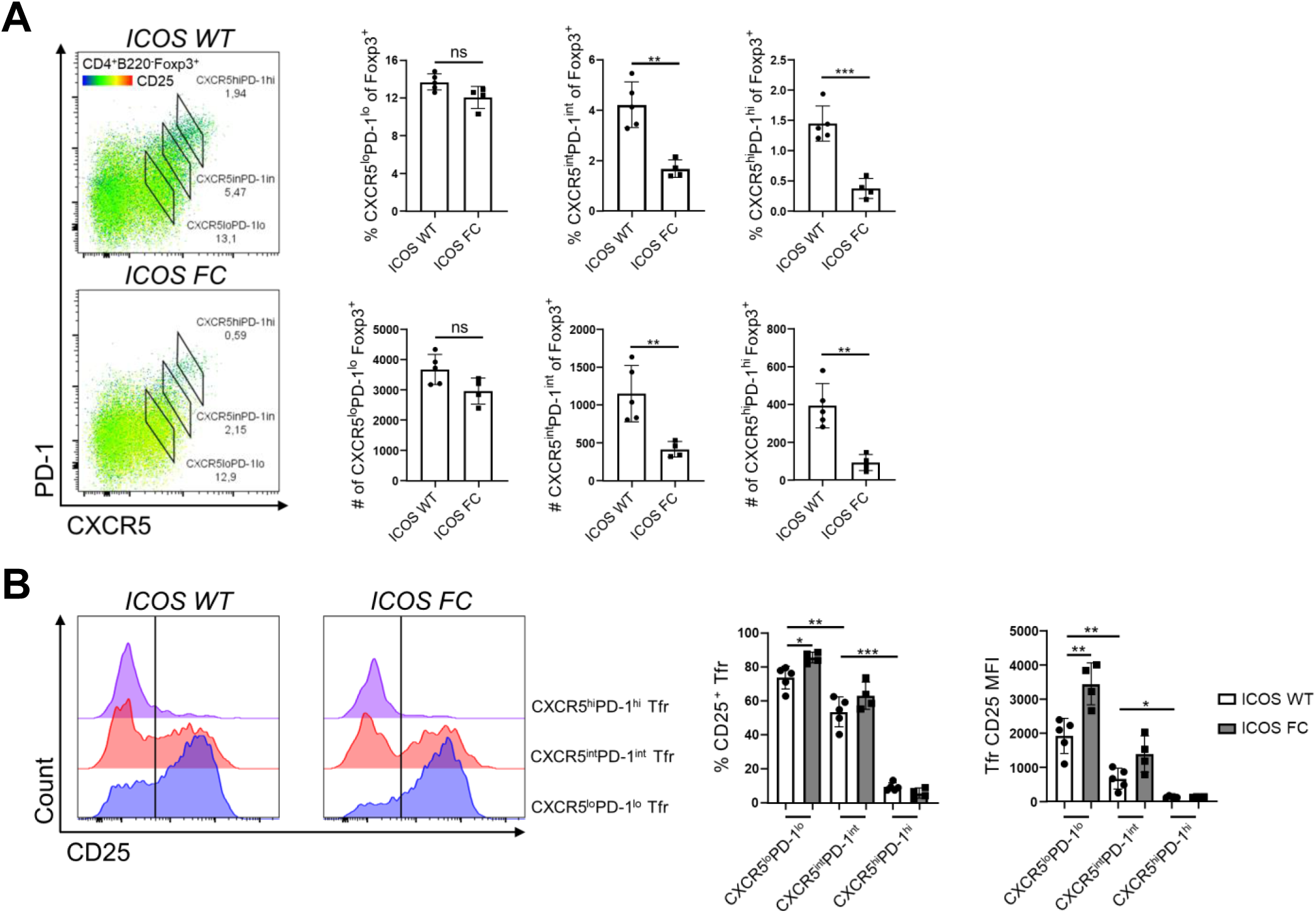
CD25 is downregulated in CXCR5hi PD-1hi GC Tfr cells. (A) Splenocytes from ICOS WT (n=5) and ICOS FC (n=4) mice were analyzed 30 dpi with IAV and Tfr subsets were defined by their relative expression levels of CXCR5 and PD-1. (B) CD25 expression levels in the Tfr subsets defined in (A) were analyzed by flow cytometry. The black bar in the histograms defines CD25^-^ and CD25^+^ populations. Data shown as mean ± SEM, \**p*<0.05, \*\**p*<0.01, \*\*\**p*<0.001. All data are representative of two independent experiments.

**Supplementary Table 1.**
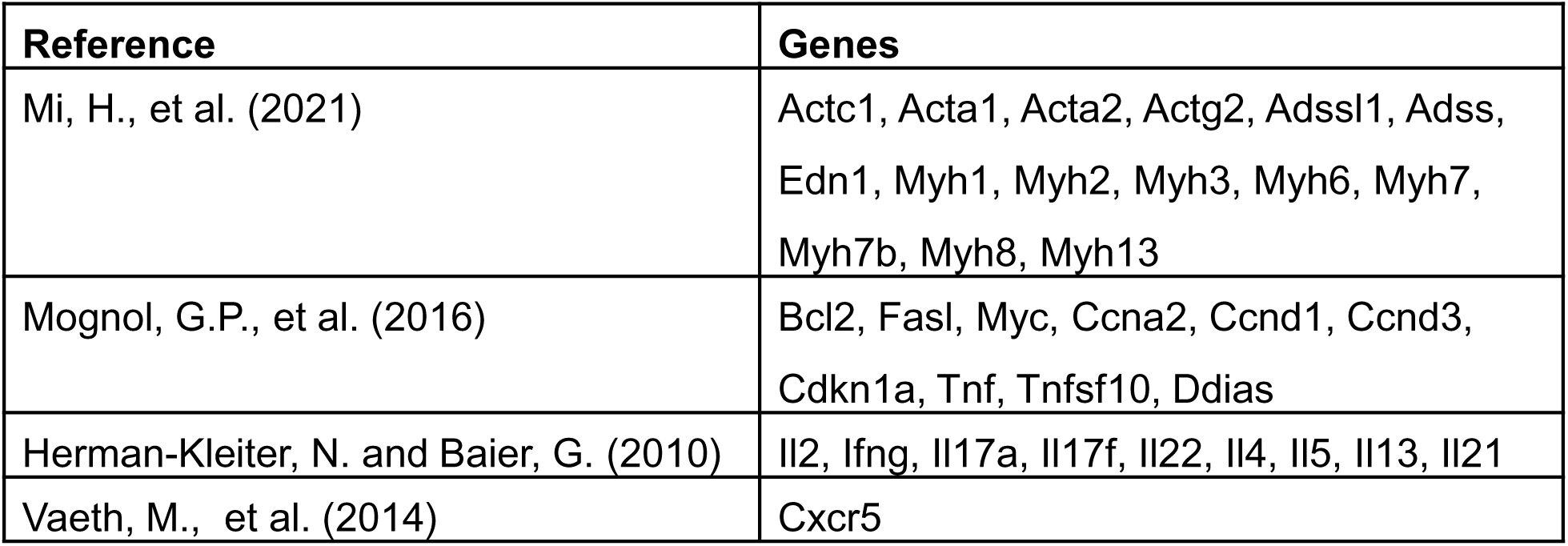
List of NFAT target genes

## Notes

### Competing Interest Statement

The authors have declared no competing interest.

### Summary of Updates

Figures 6 and 7 were removed to make the overall project more cohesive. Supplementary figure 2 was modified. Text was changed to fit the updated figures.

